# Engineered reversal of function in glycolytic yeast promoters

**DOI:** 10.1101/530717

**Authors:** Arun S. Rajkumar, Emre Özdemir, Alicia V. Lis, Konstantin Schneider, Michael K. Jensen, Jay D. Keasling

## Abstract

Promoters are key components of cell factory design, allowing precise expression of genes in a heterologous pathway. Several commonly-used promoters in yeast cell factories belong to glycolytic genes, highly expressed in actively-growing yeast when glucose is used as a carbon source. However, their expression can be suboptimal when alternate carbon sources are used, or if there is a need to decouple growth from production. Hence, there is a need for alternate promoters for different carbon sources and production schemes. In this work, we demonstrate a reversal of regulatory function in two glycolytic yeast promoters by replacing glycolytic regulatory elements with ones induced by the diauxic shift. We observe a shift in induction from glucose-rich to glucose-poor medium without loss of regulatory activity, and strong ethanol induction. Applications of these promoters were validated for expression of the vanillin biosynthetic pathway, reaching production of vanillin comparable to pathway designs using strong constitutive promoters.

---

The budding yeast *Saccharomyces cerevisiae* is a popular cell factory platform, with several genetic and molecular tools available to facilitate the production of compounds of interest for the chemical and pharmaceutical industry. The synthesis of increasingly complex molecules in turn require successful construction of large and complex heterologous pathways^1–3^, which impose an added metabolic burden by competing for required cellular resources with native metabolism. Competition for a common pool of ATP, cofactors, coenzymes and precursors between native and heterologous pathways can adversely affect growth, lengthen fermentation and decrease product titers and yields^4–6^. One way to avoid this problem is to activate the production pathway after the major growth phase. For a typical process with a fermentable carbon source, a good regulatory trigger would be the diauxic shift following fermentation^7,8^. However, most promoters commonly used for heterologous pathway construction are glycolytic (*PGK1pr, TPI1pr* or *TDH3pr*) or constitutive (*TEF1pr*), and lack the desired induction properties. Native yeast promoters have been identified for gene expression under these conditions and used for production^7–11^, yet it is unclear if sufficient native promoters with suitable properties exist to be used for large biosynthetic pathways. Therefore, using synthetic promoters with optimal regulatory output at the diauxic shift would be beneficial for cell factory design. In previous work, we developed a promoter engineering workflow to engineer yeast promoters responsive to any environmental condition given transcriptome or transcription factor binding site (TFBS) data, and functional genomics for the condition of interest if available^12^. Having used this workflow to make promoters inducible by low extracellular pH, we now use the same workflow to design promoters inducible by glucose starvation and alternating carbon sources.

Using *S. cerevisiae* CEN.PK 113-7D transcriptome data generated during the lag (10% glucose consumption), mid-exponential (75% glucose consumed), and post-exponential phases (>99% glucose consumed), we identified genes with the appropriate expression profile: strong induction upon glucose depletion. (Figure 1A). Induction under these conditions involves transcriptional activators like Cat8, Sip4, Rds2 and Adr1 (Table 1), clustered in carbon source response elements (CSREs) in promoters of interest^13–17^.Therefore, CSREs are relevant regulatory elements to ensure that promoters are activated at the diauxic shift and remain so past it. Transcriptome analysis revealed 4418 differentially expressed genes (DEGs) from the 3-way comparison, with the majority being induced at, or following, the diauxic shift (14h vs 26h), including 17 out of 19 genes (excl. *ENO2* and *PYC2*) associated with gluconeogenesis (term GO:0006094)(Figure 1B-C, Supplementary Table S1).

**Figure 1.**
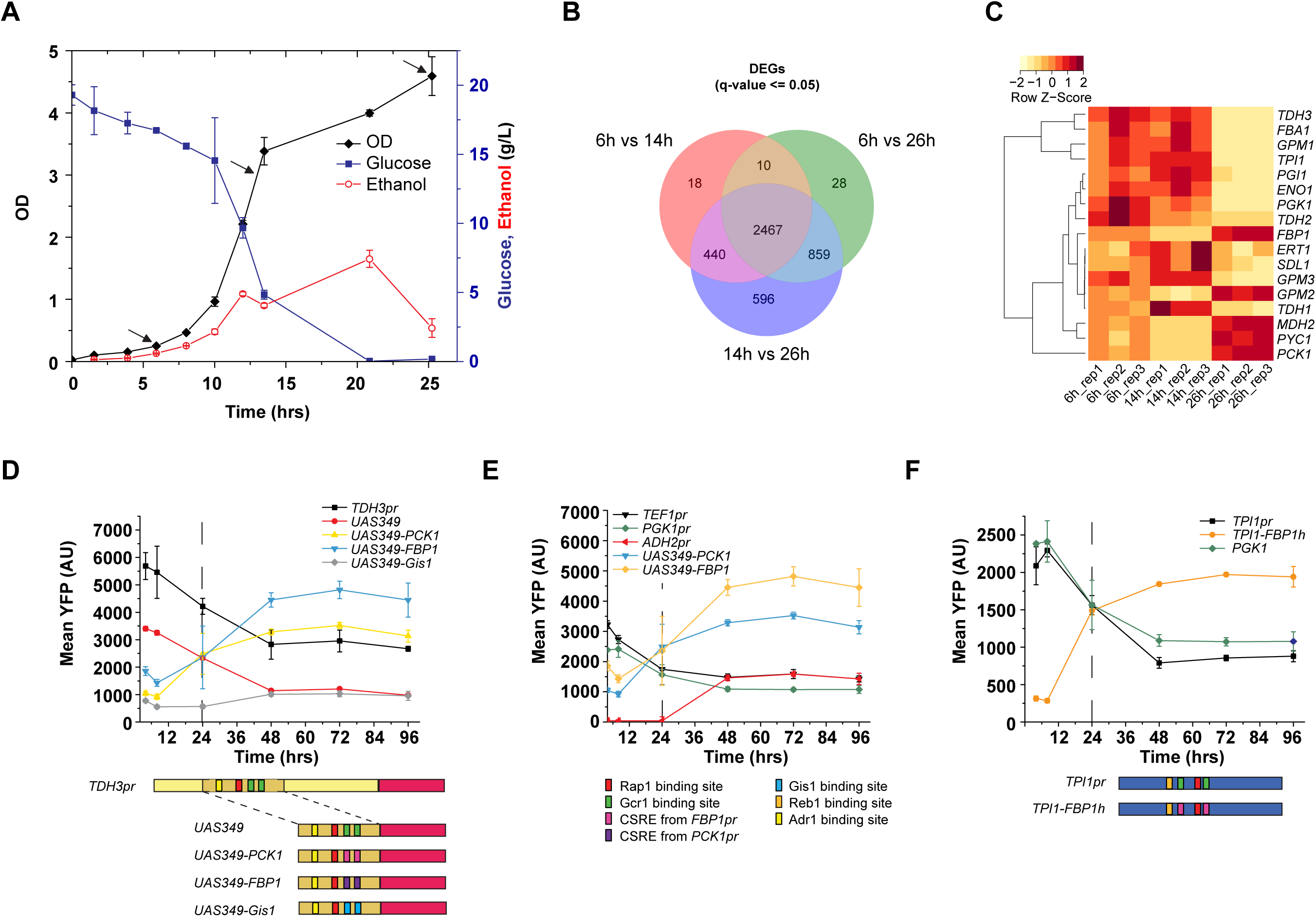
Yeast transcriptome analysis during lag, mid-exponential, and post-exponential phases, and engineering of diauxic responses in glycolytic promoters. (**A**) Growth profile of *S. cerevisiae* CEN.PK grown in Delft medium (black), and the corresponding glucose (dark blue) and ethanol (red) concentrations. Black arrows indicate sampling time-points (6h, 14h, and 26h) for RNAseq analysis. Measurements are mean +/ SEM from three biological replicates. (**B**) Venn diagram showing the numbers of significantly differentially expressed genes (DEGs) from the 3-way comparison between sampling points as indicated in (A). (**C**) Heatmap of triplicate expression profiles of 17 DEGs following the diauxic shift (14h vs 26h) under GO term for gluconeogenesis (GO:0006094). (**D**) A 200bp UAS from *TDH3pr* was fused upstream of the *TDH3* core promoter, creating *UAS-349*, a relatively strong promoter exhibiting the same trends as intact *TDH3pr* with its output decreasing over time once glucose is depleted. Replacing its Gcr1 binding sites with CSREs from gluconeogenic genes (*UAS349-PCK1, UAS349-FBP1*) reverses the induction pattern, with an output stronger than the native *TDH3pr* once glucose is depleted. Replacing Gcr1 sites with Gis1 sites (*UAS349-*Gis1) does not have the same effect. (**E**) The synthetic gluconeogenic promoters are stronger than *ADH2pr*, a native promoter strongly derepressed on glucose starvation, and the commonly-used *TEF1pr* and *PGK1pr*. (**F**) Replacing Gcr1 sites in *TPI1p* with CSREs from *FBP1pr* into *TPI1pr* cause reversal of glucose-dependent induction, even though Gcr1 sites in *TPI1pr* are differently spaced and oriented. For D-F, the dashed line indicates the time point where glucose was no longer detected as described in the Methods. All measurements in D-F represents mean +/-SEM from three biological replicates. AU = arbitrary units. At the bottom of panel D-F, illustrations outline the schematics of the promoter engineering. The sequences of the binding sites are listed in Supplementary Table S6, and all promoter sequences are listed in Supplementary Table S7.

**Table 1.** Genes associated with carbon metabolism upregulated between the log phase and diauxic shift.

| Gene | Fold-induction | Binding sites in CSREs | References for binding sites |
| --- | --- | --- | --- |
| **Annotated 'gluconeogenesis' by Gene Ontology** |  |  |  |
| *PCK1* | 472.97 | Adr1, Cat8, Sip4, Rds2 | 14,16,18,19 |
| *FBP1* | 52.67 | Adr1, Cat8, Rds2 | 14,16,19 |
| *MDH2* | 6.55 | Adr1, Cat8, Rds2 | 14,16,18 |
| *PYC1* | 1.87 | Cat8 | 14,16 |
| *GPM2* | 1.74 | None |  |
| *PYC2* | 1.46 | None |  |
| **Annotated 'carbon metabolic process' by GO_slim** |  |  |  |
| *ICL1* | 1458.62 | Cat8, Sip4 | 14,15,16,18 |
| *MLS1* | 650.68 | Adr1, Cat8, Rds2 | 14,16,19 |
| *PCK1* | 472.97 | Adr1, Cat8, Sip4,<br>Rds2 | 14,16,18,19 |
| *YIG1* | 129.24 | Adr1,Cat8 | 18 |
| *FBP1* | 52.67 | Adr1, Cat8, Rds2 | 14,16,19 |
| *MAL12* | 28.48 | Cat8 | 18 |
| *MAL11* | 29.81 | Cat8 | 18 |
| *CAT8* | 22.31 | None |  |
| *MAL32* | 16.89 | None |  |
| *GAL3* | 13.67 | Cat8 | 18 |
| *MDH2* | 6.55 | Adr1, Cat8, Rds2 | 14,16,18 |
| *DAK2* | 6.67 | None |  |
| *PYC1* | 1.87 | Cat8 | 14,16 |
| **Fold-induction of *ADH2*** |  |  |  |
| ***ADH2*** | 10.92864 | Cat8, Sip4 | 14,15,17 |

Next, we scanned the promoters for clustered TFBSs in CSREs. However, reported sequences for their binding sites contain palindromic or inverted repeats as well as stretches of ambiguous bases. For further confirmation we analyzed the discovered TFBSs for overlap with CSREs reported in the literature (Table 1; Supplementary Table S2). From this analysis, we selected experimentally validated CSREs from *PCK1* and *FBP1* as candidate elements for synthetic promoter engineering as these were annotated for gluconeogenesis (Table 1), ensuring that they would stay activated past the diauxic shift^16–18^.

These CSREs were used to engineer a 200bp upstream activating sequence (UAS) from the glycolytic *TDH3* promoter (*TDH3pr*). We chose this UAS to test the reversal of regulatory function by TFBS exchange, as its main regulatory elements are well-annotated (Figure 1D-E)^19^. YFP reporter assays revealed that the UAS fused to the *TDH3* core promoter (*UAS349*) retained 60% of *TDH3pr*’s activity, and that its Gcr1 sites were essential to promoter activity (Supplementary Figure S1). Replacing both Gcr1 sites and surrounding sequence with CSREs from either *PCK1* or *FBP1* promoters (*PCK1pr, FBP1pr*) to yield promoters *UAS349-PCK1, UAS349-FBP1* (Figure 1D) did indeed shift the induction trigger of the UAS from glucose-rich to glucose-depleted medium, with induction of *UAS349-PCK1* and *UAS349-FBP1* triggered by glucose depletion, after ~24hrs of culture. Importantly, both promoters had a stronger output than not only *UAS349* and but also *TDH3pr* - one of the strongest yeast promoters in use - which was sustained as long as glucose was not replenished. The replacement of Gcr1 sites with CSREs was sufficient and necessary for the induction. When they were exchanged with Gis1 (a TF implicated in upregulating genes at the stationary phase) binding sites instead (*UAS349-Gis1*)^20^, the promoter did not induce upon glucose starvation and remained weak throughout the assay period (Figure 1D). Ultimately, *UAS349-PCK1* and *UAS349-FBP1* were up to 3-fold stronger than other commonly-used constitutive promoters over long time periods (Figure 1E). A stronger switch in induction was achieved when the Gcr1 site exchange was made for the triose phosphate isomerase promoter (*TPI1pr*) with CSREs from *FBP1pr* (*TPI1-FBP1h*, Figure 1F, and Supplementary Figure S1)^21^.. Though the orientation, spacing and sequence context of Gcr1 sites in both *TDH3pr* and *TPI1pr* were different, we observed near-identical induction behavior. It is likely the change in induction is independent of the context of Gcr1 sites, and that their substitution could be applied to engineer induction-switching in any glycolytic promoter.

To demonstrate the usefulness of our synthetic promoters we used them to design a yeast cell factory for vanillin-β-glucoside biosynthesis (Figure 2A)^22^. The choice of product was motivated by the fact that actual vanillin-β-glucoside synthesis when using glycolytic promoters takes place in the ethanol phase ^23^. In existing cell factories and their fermentations, ~70% of the carbon ends up as toxic intermediates like protocatechuic acid (PCA), continuously produced through both glucose and ethanol phases^24,25^. We used *UAS349-PCK1* and *UAS349-FBP1* instead of constitutive promoters only for the enzymes that convert PCA to vanillin (ACAR and EntD, Figure 2A), expecting improved PCA conversion, faster growth, and ultimately similar or higher vanillin-β-glucoside production (Figure 2A). Compared to the existing pathway design relying on the use of constitutive (*TEF1pr*) and glycolytic (*PGK1pr*) promoters (C-VG) alone^23^, the use of synthetic gluconeogenic promoters to control expression of the vanillin-β-glucoside biosynthetic pathway (strains ScASR.V001 and ScASR.V003) supported production of vanillin-β-glucoside to similar levels as the C-VG strain following a 65h fermentation (Figure 2B). Likewise, the growth rate was modestly increased, and conversion of PCA towards protocatechuic aldehyde (PAL) was also improved in the engineered strains compared to C-VG (Figure 2C-D). Ultimately, ScASR.V003 produced a higher amount of vanillin-β-glucoside as a percentage of the total heterologous metabolites, suggesting that the CSRE from *FBP1pr* has the best performance for engineering gluconeogenic promoters for practical applications. We can envisage further improving PCA conversion by placing the enzyme synthesizing it from 3-dehydroshikimate, 3-DSD, under the control of another synthetic promoter responsive to glucose depletion. In this way, the entire pathway would only be activated when sufficient biomass has accumulated and PCA is given no time to accumulate and be secreted as seen in the strain designs tested here. Testing different CSREs in synthetic promoters would also allow us to select more promoters for every gene in the pathway in future configurations.

**Figure 2.**
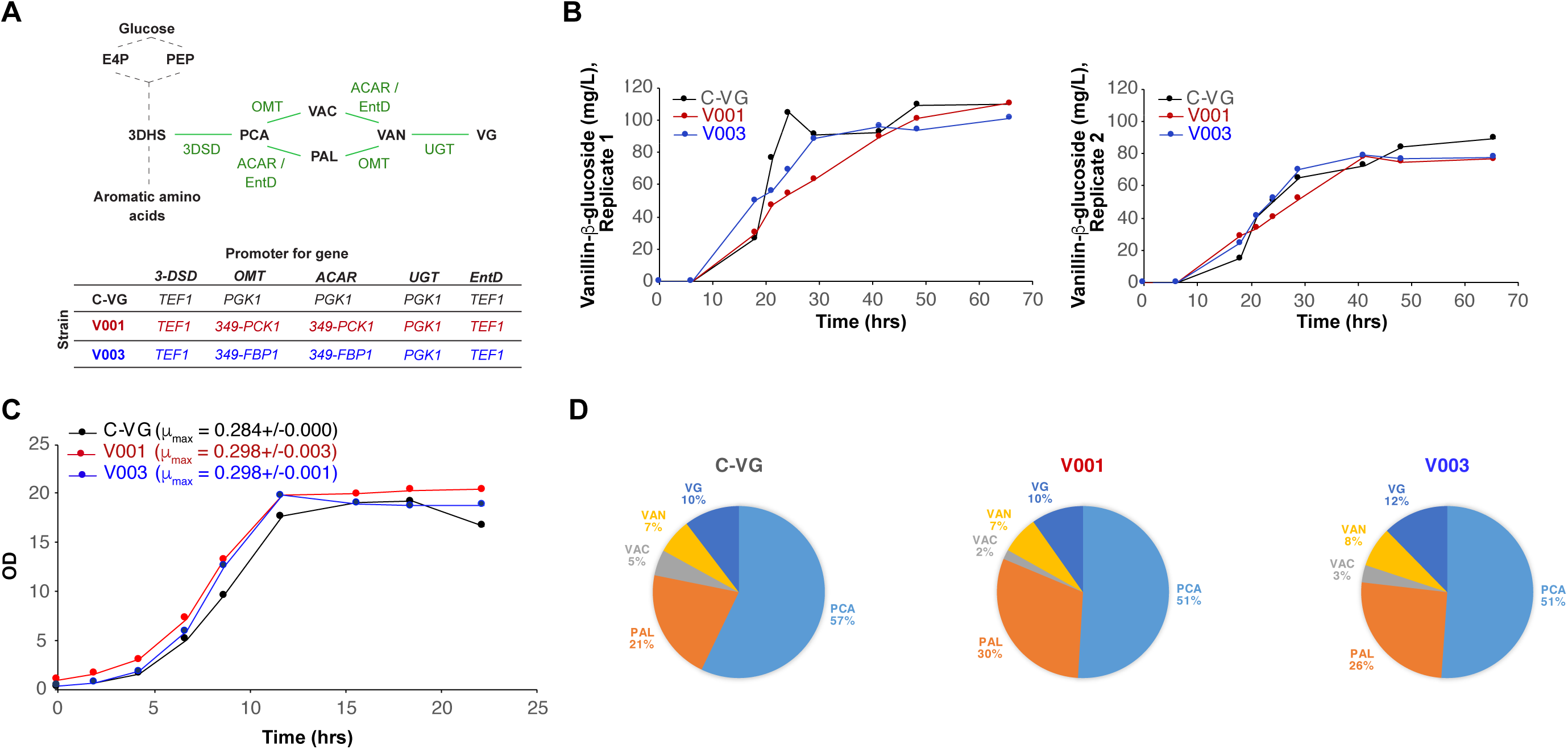
Benchmarking vanillin-β-glucoside production in yeast using synthetic glyconeogenic promoters. (**A**) Vanillin-β-glucoside pathway overview (*left*) and layout of promoter usage in control C-VG strain ^23^, and strains ScASR.V001 and ASR.V003 engineered with using synthetic gluconeogenic promoters (*right*). Starting from the shikimate pathway, the vanillin-β-glucoside pathway is marked in green. (**B**) Vanillin-β-glucoside production during 65 hrs triplicate cultivations in 5 ml deep-well plates. Data represents means +/-SEM from duplicate samples. (**C**) Representative growth profiles of vanillin-β-glucoside production strains C-VG, ScASR.V001 and ScASR.V003 cultivated in duplicates in 250mL shake-flasks. (**D**) Pie charts illustrating average relative distribution of the pathway products for each strain as a percentage of the total products of the vanillin-β-glucoside pathway. The average metabolite distribution are based on two biological replicates sampled at the end of 65 hrs fermentations (Figure 2B). The abbreviations are as follows: PCA-protocatechuic acid, PAL-protocatechuic aldelyde, VAC-vanillic acid, VAN-vanillin and VG-vanillin-β-glucoside.

The promoters engineered and put to use in this study reaffirm our promoter engineering strategy^12^ which does not require altered transcription factor expression, or orthogonal regulatory systems^26,27^. We can foresee such synthetic promoters being designed as needed, or used off-the-shelf in other applications where the bulk of products are made during ethanol consumption phases, when a product or enzyme is toxic to the cell and decoupling production from growth is potentially beneficial, or where multiple carbon sources are being used^23,28,29^. The last application is of interest for our promoters, as they also show strong induction when grown in ethanol (Supplementary Figure S2). As promoters for cell factories may not require induction at the diauxic phase, future engineering efforts will focus on finding an optimal balance of glycolytic, diauxic and gluconeogenic transcription factors to achieve constitutive high expression levels ultimately supporting the rational design of next-generation, high-performance yeast cell factories.

## MATERIALS AND METHODS

### Reporter strain construction

*S. cerevisiae* CEN.PK 113-7D was used as background strain for all experiments. Promoters were amplified from genomic DNA or gBlocks (IDT) using Phusion Master Mix with HF buffer (ThermoFisher). YFP and resistance markers were amplified from plasmid pASR0130^12^. YFP expression cassettes with different promoters were cloned into the EasyClone site XII-4 using *in vivo* assembly by homologous recombination. In brief, 500pmol of the relevant promoter, YFP, the VPS13 terminator, and *kanMX* cassette for G418 resistance marker were transformed into yeast with 500bp of homology to XII-4 on either end by the lithium acetate method onto YPD agar with 350ng/µL G418. Parts used in assembly had 50bp homology with adjacent parts added by PCR. G418-resistant colonies were screened by multiplex colony PCR as previously described^30^ checking for integration and assembly at the correct locus. Colonies with sequence-verified reporter constructs were streaked out on YPD agar, regrown in YPD with 200ng/µL G418 and preserved as glycerol stocks. All primers used for strain construction are listed in Supplementary Table S3, and all constructed strains are listed in Supplementary Table S4.

### YFP reporter assays

Reporter strains, as well as the background strain, were grown overnight in synthetic drop-out medium minus leucine (SD-leu, Sigma) containing 1.1g/L monosodium glutamate as a carbon source and 200 ng/µL G418 where appropriate. These were diluted to an OD of ~0.02 in minimal Delft medium^31^, pH 6 and grown at 30°C in deep-well microtiter plates with 300rpm agitation. The culture was sampled at 4, 8, 24, 48, 72, and 96h for measuring YFP fluorescence by flow cytometry. Cells were suitably diluted in phosphate-buffered saline (PBS, Life Technologies) and their fluorescence measured on an LSRFortessa flow cytometer (BD Biosciences) equipped with an HTS module for sampling. YFP fluorescence was measured with 488nm excitation and 530nm emission. A total of 10,000 events per sample were acquired using FACSDiva software and the resulting FCS files were analyzed using the FCSExtract utility (http://research.stowers.org/mcm/efg/ScientificSoftware/Utility/FCSExtract/index.htm), R scripts and Origin 9.1 (OriginLab) to extract the mean population fluorescence. Alternatively, the cells sampled at the same time points and diluted in PBS had their OD and YFP fluorescence (488nm excitation and 527nm emission) measured on a SynergyMX microtiter plate reader using clear-bottomed black microtiter plates (Thermofisher), and fluorescence values normalized to sample OD following the subtraction of background fluorescence from CEN.PK 113-7D. A qualitative estimate of residual glucose was determined using test strips (VWR) at each sampling point.

### Vanillin-β-glucoside production strains

The five-gene vanillin-β-glucoside biosynthetic pathway (Figure 2A) was introduced using CasEMBLR^30^ into three loci: *BGL1/EXG1* and *ADH6*, to simultaneously knock these genes out and thereby avoid product degradation^22,23^, and the EasyClone site XII-5^31^. The pathway was integrated into the genome in two steps. In the first reaction, 4pmol of each part (insertion homology arms, promoters, terminators and genes (ACAR and EntD)) were co-transformed with a gRNA expression plasmid targeting *ADH6* (clonNAT resistance) into CEN.PK 113-7D already carrying a Cas9 expression plasmid (G418 resistant). Following genotyping of transformants and sequence verification of the assemblies, colonies with correctly assembled and integrated expression cassettes were cured for loss of the *ADH6* plasmid and subsequently transformed with a gRNA plasmid targeting *EXG1* and site XII-5 along with parts for expression cassettes of OMT, UGT and 3-DSD. Colonies were screened as before, and those containing all five genes of the pathway successfully integrated and assembled were retained for fermentations. Supplementary Figure S3 gives the layout of the pathway integration and the genotyping assays. The gRNA plasmids are listed in Supplementary Table S5.

### Bioreactor cultures and HPLC analysis of metabolites

For the transcriptome analysis an overnight culture of log-phase *S. cerevisiae* grown in Delft medium was inoculated into a 500mL Delft medium at a starting OD of 0.03. The cultures were carried out in 1L Biostat Q bioreactors (Sartorius) in triplicate at 30 degrees with 800rpm agitation, with controlled aeration and pH maintained at 6 using 2M NaOH. Fermentation broth was sampled every 2 hours for the monitoring of OD, and glucose and yeast metabolites by HPLC. For the latter, the broth sample was centrifuged at 10,000g for 2 minutes, the supernatant syringe-filtered using a 0.22µm syringe filter. These were diluted two-or five-fold for analysis HPLC (UltiMate 3000, Dionex) as previously described^32^, and data acquired and analyzed using Chromeleon (Dionex/Thermofisher). For the cultivations of the vanillin-β-glucoside production strains, a single colony of each of the vanillin-β-glucoside production strains C-VG, ScASR.V001 and ScASR.V003 were picked from a YPD plate to inoculate a culture in Delft medium. Overnight cultures were used as inoculum to start duplicate cultures of each strain in Delft medium at pH 6.0 in 250mL shake-flasks with a starting OD_600nm_ of approx. 0.2. Growth rates were calculated based on OD_600nm_ measurements in the exponential growth phase. For the metabolite profile of vanillin-β-glucoside and precursors, the strains were cultivated in triplicates in 24-deepwell plates using 5mL Delft medium at pH 6.0. The respective cultures were inoculated at a starting OD_600nm_ of approx. 0.04 with a previously prepared overnight culture, and OD_600nm_ measured during the course of the cultivation by means of plate reader (SynergyMX). Samples taken at regular time intervals were extracted and quantified by HPLC as previously described^23^.

### RNAseq and transcriptome data analysis

Thirty OD units of bioreactor yeast culture grown in 6, 14 and 26h were harvested for total RNA extraction, the time points corresponding to at lag/early-log phase, mid-log phase and late-log phase/diauxic shift. Sampling points were determined from trends in growth curves. Cells were harvested from fermentation broth in chilled 15mL centrifuge tubes half-filled with ice, and following this total RNA was extracted as previously described using the RNeasy Mini kit (Qiagen)^33^. RNA concentration and quality control were performed using a Qubit 2.0 fluorometer (Life Technologies) and an Agilent 2100 Bioanalyzer with RNA 6000 Nano kit respectively (Agilent). RNAseq was performed as previously described^34^. TopHat (2.0.14) and the Cufflinks (2.2.1) suite were employed for RNAseq analysis as described previously^35^, using the reference genome and annotations for *S. cerevisiae* S288C (NCBI RefSeq GCF_000146045.2). Three biological replicates were used to determine expression levels (Fragments Per Kilobase of exon per Million fragments mapped; FPKM) for each condition. Upper-quartile normalization was preferred and reads mapping to rRNA genes were masked. Cuffdiff is used to obtain fold change differences and to perform statistical testing. A *q*-value cutoff of <0.05 was used to identify genes that have significant differential expression (Supplementary Table S1). GO and GO_slim mappings were retrieved from Saccharomyces Genome Database (http://www.yeastgenome.org). When not possible to extract due to chromosome ends, the largest possible sequence fitting the criteria was used. For transcription factor binding site analysis of CSRE, sequences spanning −700 to −125bp upstream from CDS features (or largest possible sequence fitting the criteria) were extracted and patterns matching those listed in Supplementary Table S2 were identified using biopython (Version 1.68)

### Database for RNA-seq data

RNA-seq data have been deposited in the ArrayExpress database at EMBL-EBI (www.ebi.ac.uk/arrayexpress) under accession number E-MTAB-7657 (see also Supplementary Table S1).

## ASSOCIATED CONTENT

**Supporting information.** Additional figures, tables and sequences.

## AUTHOR INFORMATION

## ACKNOWLEDGEMENTS

This work was funded by the Novo Nordisk Foundation and 7^th^ Framework program under the European Union (BioREFINE-2G, Project no. FP7-613771). We wish to thank Anna Koza for technical assistance with RNAseq, Signe Stentoft for help with strain construction, Lars Boje Petersen and Charlotte Brochner Olsen for help with HPLC, and Inger Rosenstadt for help with the bioreactor culture.

