## Supplementary Information for "Engineered reversal of function in glycolytic yeast promoters"

**Supporting Information**

Arun S. Rajkumar^1#^, Emre Özdemir^1^, Alicia V. Lis^1^, Konstantin Schneider^1^, Michael K. Jensen^1*^, and Jay D. Keasling^1-5^

^1^ Novo Nordisk Foundation Center for Biosustainability, Technical University of Denmark, Kgs. Lyngby, Denmark

^2^ Joint BioEnergy Institute, Emeryville, CA, USA

^3^ Biological Systems and Engineering Division, Lawrence Berkeley National Laboratory, Berkeley, CA, USA

^4^ Department of Chemical and Biomolecular Engineering & Department of Bioengineering, University of California, Berkeley, CA, USA

^5^ Center for Synthetic Biochemistry, Institute for Synthetic Biology, Shenzhen Institutes of Advanced Technologies, Shenzhen, China

^#^ Current address: School of Microbiology, University College Cork, Cork, Ireland.

**SUPPORTING TABLES AND FIGURES**

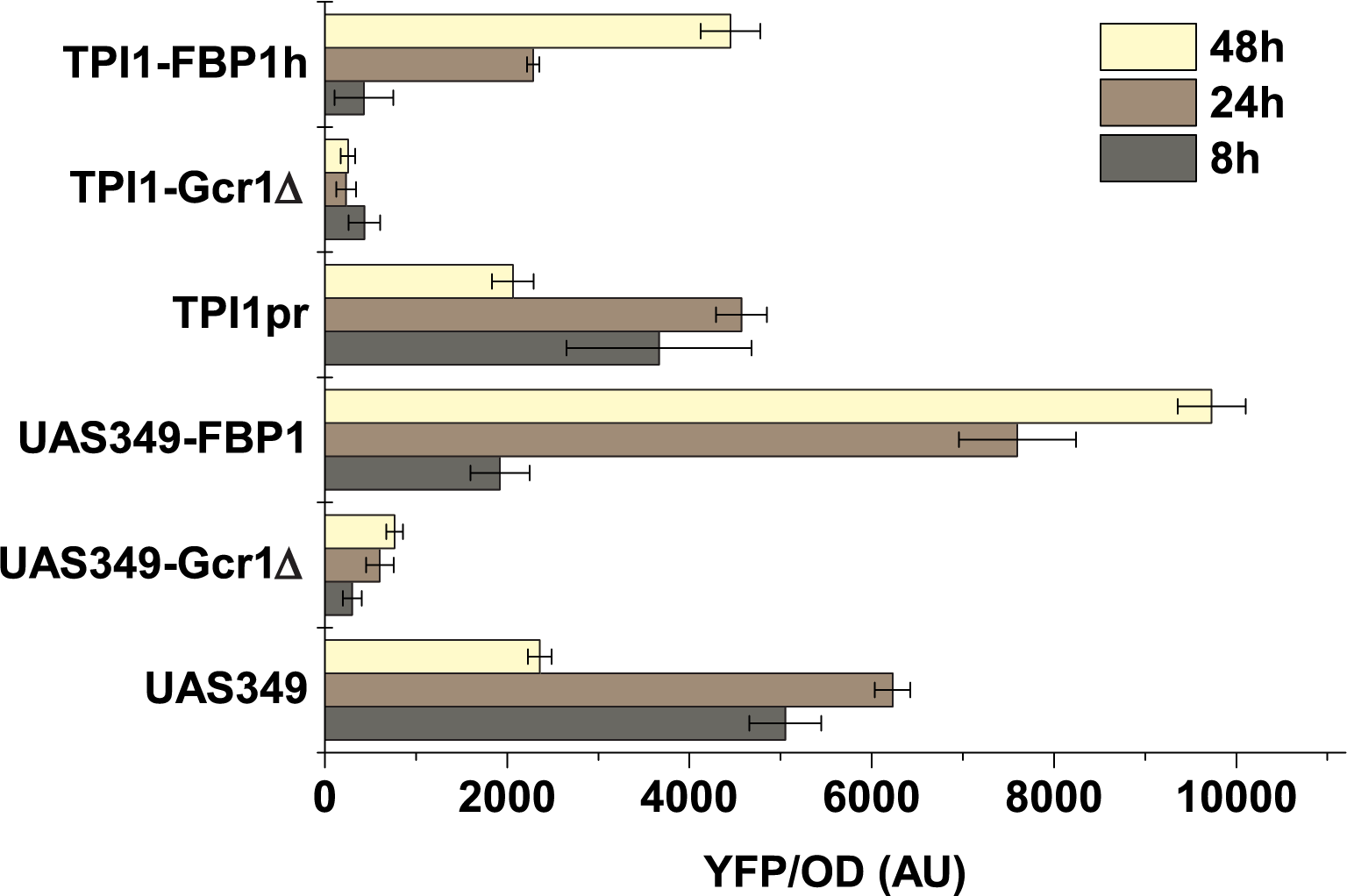

**Figure S1.** **Transcription factor requirements for glycolytic promoter function.** Removal of Gcr1 binding sites from promoters *UAS349* or *TPI1pr* (*UAS349-Gcr1D*, *TPI1pr-Gcr1D*) result in a complete loss of promoter activity as measured by YFP fluorescence normalized by OD, either in the presence (8 and 24h measurements) or absence (48h measurement) of glucose in the medium. Data from the engineered promoters *UAS349-FBP1h* and *TPI1-UAS349h* are added for comparison. Data are plotted as mean ± S.D. of 3 biological replicates.

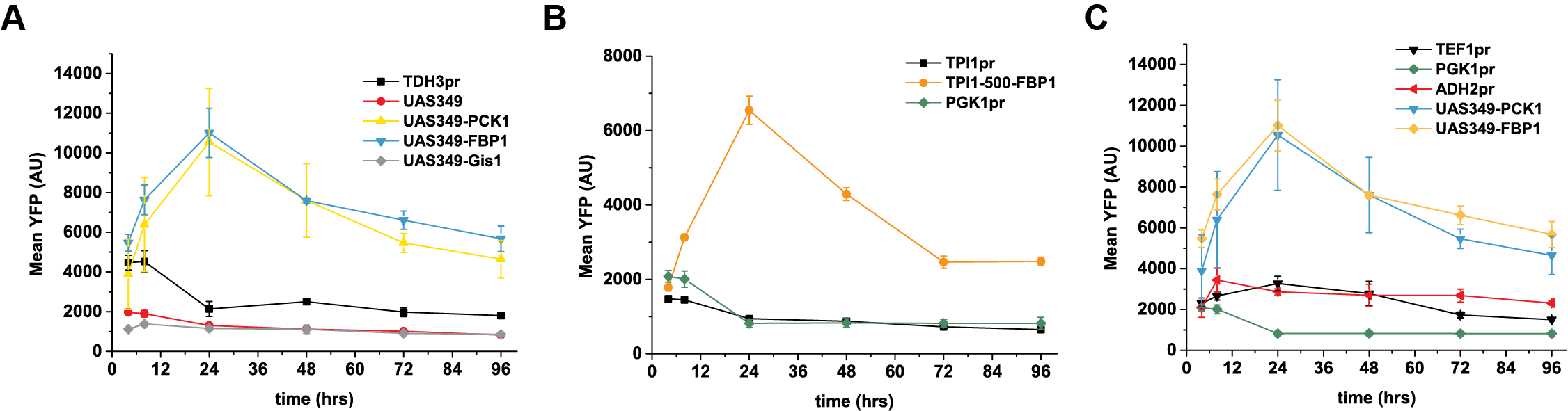

**Figure S2. Engineered promoter output with ethanol as a carbon source.** (**A**) While *TDH3pr* and *UAS349* have their output significantly reduced compared to when induced in glucose, *UAS349-PCK1h* and *UAS349-FBP1h* have a stronger output than in glucose. They also induce earlier during growth. (**B**) A similar trend is seen for *TPI1-FBP1h* versus *TPI1pr*, the three engineered promoters finally being 2-3 fold stronger than *ADH2pr* (**C**) when it is induced in ethanol. Data are plotted as mean ± S.D. of 3 biological replicates.

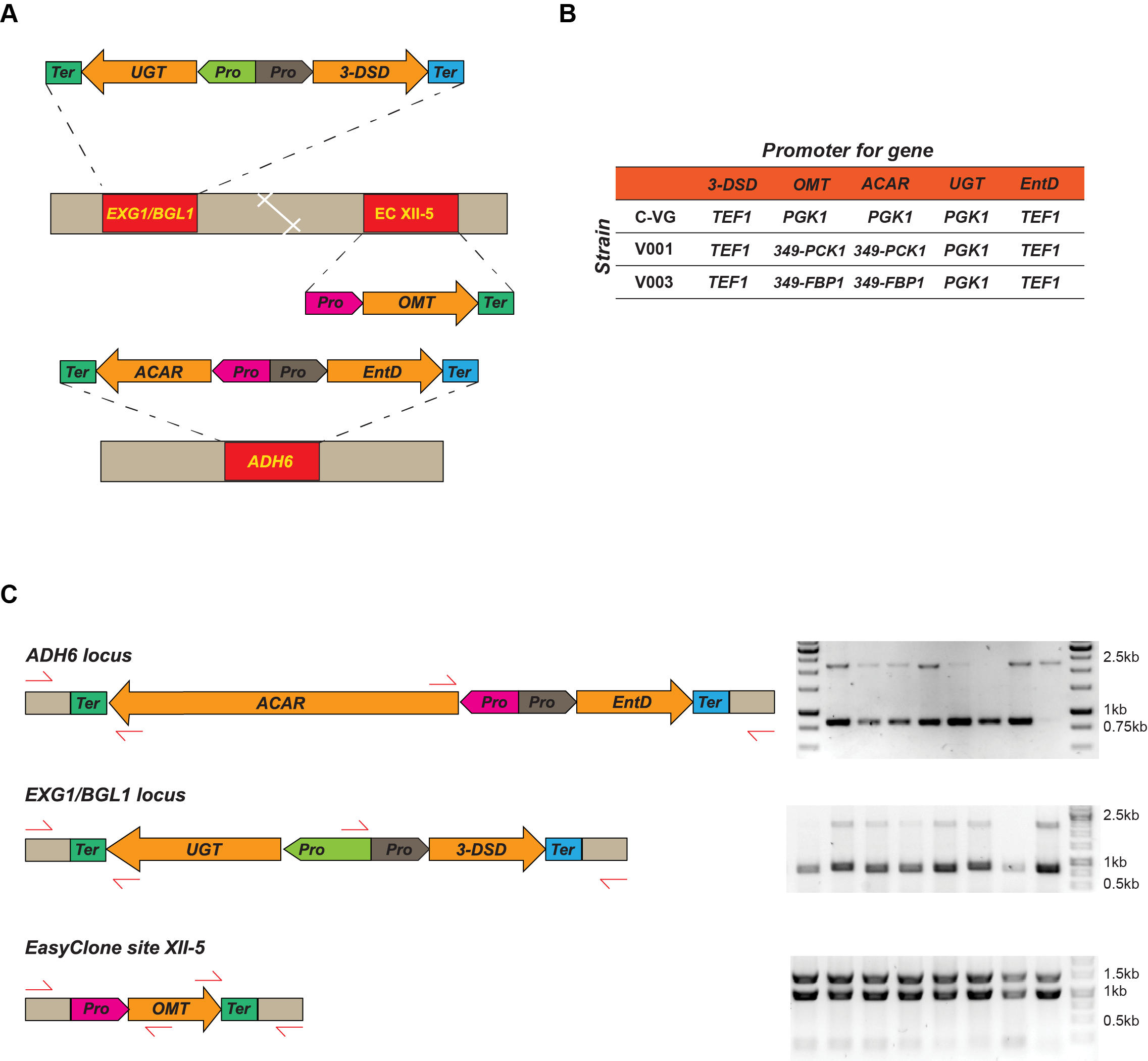

**Figure S3. Design and genotyping of vanillin-β-glucoside production strains using synthetic gluconeogenic promoters.** (**A**) Schematic outline of the design and assembly strategy for introducing the vanillin-β-glucoside biosynthesis pathway into two native genomic loci in the CEN.PK 113-7D strain background. (**B**) The combinations of genes and promoters used in the three different vanillin-β-glucoside production strains. (**C**) Multiplex genotyping strategy (*left*) for PCR validation of correct *in vivo* assembly and genomic integration of DNA parts as indicated in (A)(*right*).

**Table S1. Gene names and significance level as inferred from 3-way comparison with q-value cut-off of <0.05 to identify significantly differentially expressed genes.**

**Table S2. Transcription factor binding site occurrence in genes associated with carbon metabolism upregulated between the log phase and diauxic shift.**

| **Gene_name** | **minUAS_f** | **minUAS_rev_c** | **ADR1_f** | **ADR1_r** | **RDS2_f** | **RDS2_r** | **CAT8_f** | **CAT8_r** | **SIP4_f** | **SIP4_r** |
| --- | --- | --- | --- | --- | --- | --- | --- | --- | --- | --- |
|  | **CCNNNNNNCCG** | **CGGNNNNNNGG** | **YTGGRG** | **CYCCAR** | **YCCNTTNAKCCG** | **CGGMTNAANGGR** | **YCCNYTNRKCCG** | **CGGMYNARNGGR** | **TCCATTSRTCCGR** | **YCGGAYSAATGGA** |
| **PCK1** | CCTTTCATCCG | CGGGTGAATGG  CGGATAAAGGG | CTGGGG |  | TCCTTTCATCCG |  | TCCTTTCATCCG |  |  |  |
| **FBP1** |  | CGGACGGATGG  CGGTCGTGCGG  CGGACACCCGG |  |  |  |  |  |  |  |  |
| **MDH2** | CCTTTAATCCG  CCATTCGGCCG | CGGCATCTCGG  CGGCCAAATGG |  | CCCCAA | CCCTTTAATCCG |  | CCCTTTAATCCG  CCCATTCGGCCG | CGGCCAAATGGA |  |  |
| **PYC1** | CCGCTCCACCG |  |  |  |  |  |  |  |  |  |
| **ICL1** | CCATTCATCCG |  |  | CCCCAG | TCCATTCATCCG |  | TCCATTCATCCG |  | TCCATTCATCCGA |  |
| **MLS1** | CCATTGGGCCG  CCCCTTTCCCG  CCGGCGAGCCG | CGGCTCAATG  CGGCGAGCCGG  CGGCAGTCGGG |  | CCCCAG |  | CGGCTCAATGGA | TCCATTGGGCCG | CGGCTCAATGGA |  |  |
| **YIG1** | CCGCATTTCCG |  | TTGGGG  TTGGAG |  |  |  |  |  |  |  |
| **MAL12** | CCTCAAGCCCG |  |  |  |  |  |  |  |  |  |
| **MAL11** |  | CGGGCTTGAGG |  |  |  |  |  |  |  |  |
| **MAL32** |  |  | TTGGAG | CTCCAG |  |  |  |  |  |  |

**Table S3. Primers used in this study. Overlaps for *in vivo* assembly are in red.**

| **Name** | **Sequence** | **Function** |
| --- | --- | --- |
| XII_4_up_F | GTATCCGGCTGTTCCTTCATA | Forward XII-4 upstream homology. |
| ASR_0344_R | CTCGTGATACGCCTATTTTTATAGTGCCATAGTATGTGTGATGGA | Reverse XII-4 upstream homology, 3’ resistance marker overlap. |
| ASR_0347_F | TATTGACCACACCTCTACC$GGCATGCCAACTTTGTACTATTCCTTCCC | Forward XII-4 downstream homology, 5’ reporter construct overlap. |
| XII_4_down_R | TTTCTGCTGTACCTGGATGG | Reverse XII-4 downstream homology. |
| ASR_0307_F | AAAACAATGAGTAAAGGAGAAGAACTTTTC | Forward primer for amplification of YFP expression cassette |
| ASR_0345_R | TTTTTCCATCACACATACTATGGCACTATAAAAATAGGCGTATCACGAG | Reverse primer for YFP expression cassette, 3’ XII-4 downstream homology overlap |
| ASR_0294_F | TAAATCTGCTTTAGGTCTAGAGATCTGTTTAGCTTG | Forward primer for resistance marker |
| ASR_0346_R | CGGGGAAGGAATAGTACAAAGTTGGCATGCCGGTAGAGGTGTG | Reverse primer for resistance marker, 3’ XII-4 downstream homology overlap. |
| ASR_0318_F | AAACAGATCTCTAGACCTAAAGCAGATTTAATAAAAAACACGCTTTTTCAGTTCG | Forward primer for TDH3pr, 5’ overlap with resistance marker |
| ASR_0310_F | AAACAGATCTCTAGACCTAAAGCAGATTTATTAGTCAAAAAATTAGCCTTTTAATT | Forward primer for UAS349 and derivatives, 5’ overlap with resistance marker |
| ASR_0031_R | GTTCTTCTCCTTTACTCATTGTTTTTTTGTTTGTTTATGTGTGTTTATTCGA | Reverse primer for TDH3pr and UAS3 derivatives, 3’ overlap with YFP |
| ASR_0342_F | AAACAGATCTCTAGACCTAAAGCAGATTTAATTTAAACTGTGAGGACCTTAATACATTC | Forward primer for TPI1pr and derivatives, 5’overlap with resistance marker |
| ASR_0343_R | GTTCTTCTCCTTTACTCATTGTTTTTTTTAGTTTATGTATGTGTTTTTTGTAGT | Reverse primer for TPI1pr and derivatives, 3’ overlap with YFP |
| ASR_0273_F | AAACAGATCTCTAGACCTAAAGCAGATTTAGCACACACCATAGCTTCAAA | Forward primer for TEF1pr, 5’overlap with resistance marker |
| ASR_0274_R | GTTCTTCTCCTTTACTCATTGTTTTTTGTAATTAAAACTTAGATTAGATTGCTATG | Reverse primer for TEF1pr, 3’ overlap with YFP |
| ASR_0335_F | AAACAGATCTCTAGACCTAAAGCAGATTTAGGAAGTACCTTCAAAGAATGG | Forward primer for PGK1pr, 5’overlap with resistance marker |
| ASR_0336_R | GTTCTTCTCCTTTACTCATTGTTTTTGTTTTATATTTGTTGTAAAAAGTAGATAA | Reverse primer for PGK1pr, 3’ overlap with YFP |
| ASR_0448_F | AAACAGATCTCTAGACCTAAAGCAGATTTATCTCTCCGGTTACAGCCTG | Forward primer for ADH2pr, 5’overlap with resistance marker |
| ASR_0449_R | GTTCTTCTCCTTTACTCATTGTTTTTGTGTATTACGATATAGTTAATAGTTGATAGTT | Reverse primer for ADH2pr, 3’ overlap with YFP |
| ASR_0020_F | ACTGACGTCGAAGGCTCTTT | Forward colony PCR primer to verify insertion at XII-4, 5’ end. Primes 104bp upstream of site XII-4, and is used with ASR_0307_F. |
| XII-4-up-out-sq ID897 | CGTGAAATCTCTTTGCGGTAG | Forward colony PCR primer to verify insertion at XII-4, 3’ end. Primes 100bp downstream of XII-4, and is used with ASR_0110_F. |
| ASR_0110_F | GCCATCTAAGTTGGGACACAGATAAGCGAATTTCTTATGATTTATGATTTTTATT | Reverse colony PCR primer to verify insertion at XII-4, 3’end. |
| ASR_0288_R | CGCTTGACATCTACTATATGTAAG | Forward colony PCR primer to verify assembly of promoter-reporter cassette. Also used for sequencing. |
| ASR_0314_R | GGGATGTATGGGCTAAATGT | Reverse colony PCR primer to verify assembly of promoter-reporter cassette. Also used for sequencing. |
| ASR_0083_SEQ | TCACCATCTAATTCAACAAGAATTG | Sequencing primer for promoter. |
| ASR_V001 | ACGAGCCTGAGACAAGCC | EXG1_US_F |
| ASR_V002 | TATTGACCACACCTCTACCGGCATGTTTAGTTGGTAATTAACTAGAAAAAGAAAG | EXG1_US_R, overlap with ADH1 terminator |
| ASR_V003 | CGTGGACTTGTCACGTGGTGCCTAGGCGAATTTCTTATGATTTATGATTTTTATT | Forward primer for tADH1, overlap with UGT1 |
| ASR_V004 | TTTTTCTAGTTAATTACCAACTAAACATGCCGGTAGAGGTGTG | Reverse primer for tADH1, overlap with EXG1_US |
| ASR_V005 | TACTTTTTACAACAAATATAAAAC  AATGCATATCACAAAACCACAC | Forward primer for UGT1, overlap with PGK1pr |
| ASR_V006 | AAATCATAAATCATAAGAAATTCGCCTAGGCACCACGTGACAAGT | Reverse prime for UGT1, overlap with tADH1 |
| ASR_V007 | CGGCGTGTGGTTTTGTGATATGCATTGTTTTATATTTGTTGTAAAAAGTAGATAA | Reverse primer for PGK1, overlap with UGT1 |
| ASR_V008 | AACATTTTGAAGCTATGGTGTGTGCAAGAAGTACCTTCAAAGAATGG | Forward primer for PGK1, overlap with TEF1 |
| ASR_V009 | ACCCCATTCTTTGAAGGTACTTCTTGCACACACCATAGCTTCAAA | Forward primer for TEF1, overlap with PGK1 |
| ASR_V010 | GCTCCTTAGTGTCACCCATTGTTTTTTGTAATTAAAACTTAGATTAGATTGCTATG | Reverse primer for TEF1, overlap with HsOMT |
| ASR_V011 | ATCTAAGTTTTAATTACAAAAAACAATGGGTGACACTAAGGAGC | Forward primer for HsOMT, ovelap with TEF1 |
| ASR_V012 | TCCTTCCTTTTCGGTTAGAGCGGATTTATGGACCAGCTTCAGAAC | Reverse primer for HsOMT, overlap with tCYC1 |
| ASR_V013 | TCCAGGTTCTGAAGCTGGTCCATAAATCCGCTCTAACCGAAAAGGA | Forward primer for tCYC1, overlap with HsOMT |
| ASR_V014 | CTAAAATGAGCGGACTGAGGGCGACTTCTCAAGCAAGGTTTTCAGTATAATG | Reverse primer for tCYC1, overlap with EXG1_DS |
| ASR_V015 | ATTATACTGAAAACCTTGCTTGAGAAGTCGCCCTCAGTCCGCTC | EXG1_DS_F, overlap with tCYC1 |
| ASR_V016 | GTTGTTTAAGTTCTTTTATCCTTCCTTAGATAACA | EXG1_DS_R |
| ASR_V017 | AATACGCACACATACGCGCAT | US CPCR_F, 91bp US of EXG1_US |
| ASR_V018 | CCAAGGAGTGTCAACGGTT | US_CPCR_R,57bp US of UGT's Stop codon, product 840bp |
| ASR_V019 | GTCATCGACGTTATCTCTACTATAG | DS_CPCR_F, 105bp into PGK1 |
| ASR_V020 | ATGTTCTTTCGTTTGGATGAGG | DS_CPCR_R, 84bp DS of EXG1_DS, product 1.95kb for original config,2.4kb with DSD |
| ASR_V021 | TTCCAAGCTCGATCACCGG | EXG1_jn_F, 79 bp DS of UGT's Start codon |
| ASR_V022 | GGAGTCCGAGAAAATCTGGAAGAGTA | EXG1_jn_R,63 1126bp DS into TEF1, product for original config |
| ASR_V023 | TTTTCATTTTCAACTTGGTAATGAC | ADH6_US_F |
| ASR_V024 | TATTGACCACACCTCTACCGGCATGGATTTTGGCTTTTCTTGTTGTTG | ADH6_US_R, overlap with ADH1 terminator |
| ASR_V025 | TTTGGAATTGTTACAATTGTTATAAGCGAATTTCTTATGATTTATGATTTTTATT | Forward primer for tADH1, overlap with ACAR |
| ASR_V026 | CACAACAACAAGAAAAGCCAAAATCCATGCCGGTAGAGGTGTG | Reverse primer for tADH1, overlap with ADH6_US |
| ASR_V027 | TACTTTTTACAACAAATATAAAACAATGGCTGTTGATTCACCAG | Forward primer for ACAR, overlap with PGK1pr |
| ASR_V028 | AAATCATAAATCATAAGAAATTCGCTTATAACAATTGTAACAATTCCAAATC | Reverse prime for ACAR, overlap with tADH1 |
| ASR_V029 | TCTCATCTGGTGAATCAACAGCCATTGTTTTATATTTGTTGTAAAAAGTAGATAA | Reverse primer for PGK1, overlap with ACAR |
| ASR_V030 | GTAGTTTTCATATCGACCATTGTTTTTTGTAATTAAAACTTAGATTAGATTGCTATG | Reverse primer for TEF1, overlap with EntD |
| ASR_V031 | ATCTAAGTTTTAATTACAAAAAACAATGGTCGATATGAAAACTACGC | Forward primer for EntD, ovelap with TEF1 |
| ASR_V032 | TCCTTCCTTTTCGGTTAGAGCGGATTTAATCGTGTTGGCACAGC | Reverse primer for EntD, overlap with tCYC1 |
| ASR_V033 | CATAACGCTGTGCCAACACGATTAAATCCGCTCTAACCGAAAAGGA | Forward primer for tCYC1, overlap with HsOMT |
| ASR_V034 | CTACATTTATCAAGAGCTTGACAACTTCTCAAGCAAGGTTTTCAGTATAATG | Reverse primer for tCYC1, overlap with ADH6_DS |
| ASR_V035 | ATTATACTGAAAACCTTGCTTGAGAAGTTGTCAAGCTCTTGATAAATGT | ADH6_DS_F, overlap with tCYC1 |
| ASR_V036 | GATAATTGTCTCGAGCCAAA | ADH6_DS_R |
| ASR_V037 | TACCTCACCTGAGTTTTGCTT | US CPCR_F, 91bp US of ADH6_US |
| ASR_V038 | CTCCATTGATTGATAAGTATGTTTCTGA | US_CPCR_R,53bp US of ACAR's Stop codon, product 845bp |
| ASR_V039 | AGTATCAACCACTAAAGCGAAAAAC | DS_CPCR_R, 84bp DS of ADH61_DS, use with DS_CPCR_F for EXG1, product 1.92kb for original config |
| ASR_V040 | CTTGCTCATCTTCAGCAAACAATTGAG | ADH6_jn_F, 79 bp DS of ACAR's Start codon, use with EXG1_jn_R for 1.17bp in original config |
| ASR_V041 | AACACACATAAACAAACAAAAAAACAATGCATATCACAAAACCACAC | Forward primer for UGT1, overlap with TDH3 core |
| ASR_V042 | TGTGGTTTTGTGATATGCATTGTTTTTTTGTTTGTTTATGTGTGTTTATTCGA | Reverse primer for TDH3 core, overlap with UGT1 |
| ASR_V043 | AACACACATAAACAAACAAAAAAACAATGGCTGTTGATTCACCAG | Forward primer for ACAR, overlap with TDH3 core |
| ASR_V044 | TCTGGTGAATCAACAGCCATTGTTTTTTTGTTTGTTTATGTGTGTTTATTCGA | Reverse primer for TDH3 core, overlap with ACAR |
| ASR_V045 | AACATTTTGAAGCTATGGTGTGTGCTTAGTCAAAAAATTAGCCTTTTAATT | Forward primer for '349', overlap with TEF1 |
| ASR_V046 | ATTAAAAGGCTAATTTTTTGACTAAGCACACACCATAGCTTCAAA | Forward primer for TEF1, overlap with '349' |
| ASR_V047 | CAATCTGGCGGCTTGAGT | Forward primer for XII-5-up |
| ASR_V048 | AACATTTTGAAGCTATGGTGTGTGCGCTGATGTGACACTGTGACAAT | Reverse primer for XII-5-up, overlap with TEF1 promoter |
| ASR_V049 | TTTATTGTCACAGTGTCACATCAGCGCACACACCATAGCTTCAAA | Forward primer for TEF1, overlap with XII-5-up |
| ASR_V050 | ATGGCGAGTTTGGAAGGCATTGTTTTTTGTAATTAAAACTTAGATTAGATTGCTATG | Reverse primer for TEF1, overlap with 3-DSD |
| ASR_V051 | ATCTAAGTTTTAATTACAAAAAACAATGCCTTCCAAACTCGCC | Forward primer for 3-DSD, overlap with TEF1 |
| ASR_V052 | TCCTTCCTTTTCGGTTAGAGCGGATTTACAAAGCCGCTGACAGC | Reverse primer for 3-DSD, overlap with tCYC1 |
| ASR_V053 | GCTGTCGCTGTCAGCGGCTTTGTAAATCCGCTCTAACCGAAAAGGA | Forward primer for tCYC1, overlap with 3-DSD |
| ASR_V054 | GCTTGCTGTCAAACTTCTGAGTTGCTTCTCAAGCAAGGTTTTCAGTATAATG | Reverse primer for tCYC1, overlap with XII-5-down |
| ASR_V055 | TTATACTGAAAACCTTGCTTGAGAAGCAACTCAGAAGTTTGACAGC | XII-5-down_F, overlap with tCYC1 |
| ASR_V056 | CTGCGATACCTTTTGTGAT | XII-5-down_R |
| ASR_V057 | CGATCCAGAACTCGAGCT | US_CPCR_R, use with 899(137bp up of up, so 454+137), 337 bp into 3-DSD, 1354bp product, use with EXG1_jn_F for 1.8kb? |
| ASR_V058 | ACACAGCAGCCCATCAGG | DS_CPCR_F, use with 900 (676+97), 45bp upstream of 3-DSD's Stop, 991bp product |
| ASR_V063 | ACCCCATTCTTTGAAGGTACTTCTTGCTGATGTGACACTGTGACAAT | Reverse primer for XII-5-up, overlap with PGK1 promoter |
| ASR_V064 | TTTATTGTCACAGTGTCACATCAGCAAGAAGTACCTTCAAAGAATGG | Forward primer for PGK1, overlap with XII-5-up |
| ASR_V065 | TTCTTTGCTCCTTAGTGTCACCCATTGTTTTATATTTGTTGTAAAAAGTAGATAA | Reverse primer for PGK1, overlap with HsOMT |
| ASR_V066 | TACTTTTTACAACAAATATAAAACAATGGGTGACACTAAGGAGC | Forward primer for HSOMT, overlap with PGK1 |
| ASR_V067 | AAATCATAAATCATAAGAAATTCGCTTATGGACCAGCTTCAGAAC | Reverse primer for HsOMT, overlap with tADH1 |
| ASR_V068 | TCCAGGTTCTGAAGCTGGTCCATAAGCGAATTTCTTATGATTTATGATTTTTATT | Forward primer for tADH1, overlap with HsOMT |
| ASR_V069 | GCTTGCTGTCAAACTTCTGAGTTGCCATGCCGGTAGAGGTGTG | Reverse primer for tADH1, overlap with XII-5-down |
| ASR_V070 | TATTGACCACACCTCTACCGGCATGGCAACTCAGAAGTTTGACAGC | Forward primer for XII-5-down, overlap with tADH1 |
| ASR_V073 | ATTAAAAGGCTAATTTTTTGACTAAGCTGATGTGACACTGTGACAAT | Reverse primer for XII-5-up, overlap with '349' |
| ASR_V074 | AACACACATAAACAAACAAAAAAACAATGGGTGACACTAAGGAGC | Forward primer for HsOMT, overlap with TDH3 core |
| ASR_V075 | TTTATTGTCACAGTGTCACATCAGCTTAGTCAAAAAATTAGCCTTTTAATT | Forward primer for '349', overlap with XII-5-up |
| ASR_V076 | GAGTAATAGCAGCACAGTCTG | US_CPCR_R, 298 bp ds of OMT's Start codon, use with 899 (so 454+137+ 984+298)-a 1873bp product with PGK1, 1323bp with 349s |
| ASR_V077 | CCAGGTTCTGAAGCTGGTC | DS_CPCR_F, 24 bp us of OMT's Stop codon, yields a 989bp product with primer 900 |
| ASR_V078 | GCTCCTTAGTGTCACCCATTGTTTTTTTGTTTGTTTATGTGTGTTTATTCGA | Reverse primer for TDH3 core, overlap with HsOMT |
| ASR_V076_alt | GTTGTCAGCTAACAAAACAGTAC | US_CPCR_R for 349 cassette, 510bp into OMT's Start codon |
| ASR_V081 | CTAAAGCTATGTTGGGTGTTG | Primer inside ACAR, covers 1.5kb us of its Stop codon, to use to amplify gene for sequencing |
| ASR_V082 | CTCATGTAACAAGTTAGAGAAAGAC | Primer approx 2.1kb ds of ACAR's Start Codon, use to amplify gene for sequencing |
| ASR_V083 | CCACCGAAGTTGATTTGCTT |  |
| ASR_V084 | GTGGGAGTAAGGGATCCTGT |  |

**Table S4. Strain list**

| **Strain** | **Genotype** | **Reference** |
| --- | --- | --- |
| CEN.PK 113-7D | *MATa MAL 2-8^C^ SUC2* | Euroscarf |
| ScASR.0076i | CEN.PK 113-7D *XII-4(kanMX-TDH3pr-YFP-tVPS13)* | This study |
| ScASR.0077i | CEN.PK 113-7D *XII-4(kanMX-UAS349pr-YFP-tVPS13)* | This study |
| ScASR.0232i | CEN.PK 113-7D *XII-4(kanMX-UAS349-FBP1h-YFP-tVPS13)* | This study |
| ScASR.0233i | CEN.PK 113-7D *XII-4(kanMX-UAS349-PCK1h-YFP-tVPS13)* | This study |
| ScASR.0241i | CEN.PK 113-7D *XII-4(kanMX-UAS349-Gis1-YFP-tVPS13)* | This study |
| ScASR.0242i | CEN.PK 113-7D *XII-4(kanMX- UAS349-Gcr1D*,*-YFP-tVPS13)* | This study |
| ScASR.0131i | CEN.PK 113-7D *XII-4(kanMX-TPI1pr-YFP-tVPS13)* | This study |
| ScASR.0236i | CEN.PK 113-7D *XII-4(kanMX-TPI1-FBP1h -YFP-tVPS13)* | This study |
| ScASR.0245i | CEN.PK 113-7D *XII-4(kanMX- TPI1-Gcr1D-YFP-tVPS13)* | This study |
| ScASR.0094i | CEN.PK 113-7D *XII-4(kanMX-TEF1pr-YFP-tVPS13)* | This study |
| ScASR.0125i | CEN.PK 113-7D *XII-4(kanMX-PGK1pr-YFP-tVPS13)* | This study |
| ScASR.0223i | CEN.PK 113-7D *XII-4(kanMX-ADH2pr-YFP-tVPS13)* | This study |
| C-VG | *MATa MAL 2-8^C^ SUC2 XII-2(pTEF1-HsOMT,pPGK1-UGT)XII1(pTEF1-PPT,pPGK1-ACAR)XII-5(pTEF1-3DSD)*  *Δbgl1::loxP Δadh6::KanMX* | Strucko et al ^1^ |
| ScASR.V001 | CEN.PK 113-7D *ADH6(349-PCK1-ACAR-tADH1 TEF1pr-EntD-tCYC1) EXG1(PGK1pr-UGT-tADH1 TEF1pr-3-DSD-tCYC1) XII-5 (349-PCK1-OMT-tADH1)* | This study |
| ScASR.V003 | CEN.PK 113-7D *ADH6(349-FBP1-ACAR-tADH1 TEF1pr-EntD-tCYC1) EXG1(PGK1pr-UGT-tADH1 TEF1pr-3-DSD-tCYC1) XII-5 (349-FBP1-OMT-tADH1)* | This study |

**Table S5. Plasmid list**

| **Name** | **Features** | **Reference** |
| --- | --- | --- |
| pCfB2312 | *TEF1pr-Cas9-CYC1t-kanMX*, CEN/ARS plasmid | Stovicek et al ^2^ |
| pTAJAK-71 | · pESC-NatMXsyn-USER | Jessop-Fabre et al. ^3^ |
| pASR.G0005 | pTAJAK-71 *SNR52pr-gRNA.ADH6-tSUP4*  gRNA sequence: GACATTaagatCGAAGCATG | This study |
| pASR.G0007 | pTAJAK-71 *SNR52pr-gRNA.EXG1 XII-5-tSUP4*  gRNA sequence: AGATACTacgaTTATGACCA | Strucko et al ^1^ |
| pXII1-23 | *PGK1pr-ACAR-tADH1 TEF1pr-EntD-tCYC1* | Strucko et al ^1^ |
| pXII2-54 | *PGK1pr-UGT-tADH1 TEF1pr-OMT-tCYC1* | Strucko et al ^1^ |
| pXII5-01 | *TEF1pr-3DSD-tCYC1* | Strucko et al ^1^ |

**Table S6. Transcription factor binding site motifs used for promoter engineering**

| **Site** | **Sequence** | **Reference** |
| --- | --- | --- |
| CSRE from PCK1pr | ACGGGTGAATGGAGATCTGG | Soontorngun et al ^4^ |
| CSRE from FBP1pr | CCGGACGGATGGAATCGCCG | Soontorngun et al ^4^ |
| Gis1 binding site | TAGGGAT | Zhang et al ^5^ |

**Table S7. Sequences of all synthetic promoters engineered in this study.** Engineered binding sites are marked in boldface.

| **Promoter** | **Sequence** |
| --- | --- |
| TDH3pr | GAATAAAAAACACGCTTTTTCAGTTCGAGTTTATCATTATCAATACTGCCATTTCAAAGAATACGTAAATAATTAATAGTAGTGATTTTCCTAACTTTATTTAGTCAAAAAATTAGCCTTTTAATTCTGCTGTAACCCGTACATGCCCAAAATAGGGGGCGGGTTACACAGAATATATAACATCGTAGGTGTCTGGGTGAACAGTTTATTCCTGGCATCCACTAAATATAATGGAGCCCGCTTTTTAAGCTGGCATCCAGAAAAAAAAAGAATCCCAGCACCAAAATATTGTTTTCTTCACCAACCATCAGTTCATAGGTCCATTCTCTTAGCGCAACTACAGAGAACAGGGGCACAAACAGGCAAAAAACGGGCACAACCTCAATGGAGTGATGCAACCTGCCTGGAGTAAATGATGACACAAGGCAATTGACCCACGCATGTATCTATCTCATTTTCTTACACCTTCTATTACCTTCTGCTCTCTCTGATTTGGAAAAAGCTGAAAAAAAAGGTTGAAACCAGTTCCCTGAAATTATTCCCCTACTTGACTAATAAGTATATAAAGACGGTAGGTATTGATTGTAATTCTGTAAATCTATTTCTTAAACTTCTTAAATTCTACTTTTATAGTTAGTCTTTTTTTTAGTTTTAAAACACCAAGAACTTAGTTTCGAATAAACACACATAAACAAACAAA |
| TDH3 core promoter | CTAATAAGTATATAAAGACGGTAGGTATTGATTGTAATTCTGTAAATCTATTTCTTAAACTTCTTAAATTCTACTTTTATAGTTAGTCTTTTTTTTAGTTTTAAAACACCAAGAACTTAGTTTCGAATAAACACACATAAACAAACAAA |
| UAS349pr | TTAGTCAAAAAATTAGCCTTTTAATTCTGCTGTAACCCGTACATGCCCAAAATAGGGGGCGGGTTACACAGAATATATAACATCGTAGGTGTCTGGGTGAACAGTTTATTCCTGGCATCCACTAAATATAATGGAGCCCGCTTTTTAAGCTGGCATCCAGAAAAAAAAAGAATCCCAGCACCAAAATATTGTTTTCTTCACTAATAAGTATATAAAGACGGTAGGTATTGATTGTAATTCTGTAAATCTATTTCTTAAACTTCTTAAATTCTACTTTTATAGTTAGTCTTTTTTTTAGTTTTAAAACACCAAGAACTTAGTTTCGAATAAACACACATAAACAAACAAA |
| UAS349-PCK1h | TTAGTCAAAAAATTAGCCTTTTAATTCTGCTGTAACCCGTACATGCCCAAAATAGGGGGCGGGTTACACAGAATATATAACATCGTAGGTGTCTGGGTGAACAGTTTATTCCT**ACGGGTGAATGGAGATCTGG**GAGCCCGCTTTTTA**ACGGGTGAATGGAGATCTGG**AAGAATCCCAGCACCAAAATATTGTTTTCTTCACTAATAAGTATATAAAGACGGTAGGTATTGATTGTAATTCTGTAAATCTATTTCTTAAACTTCTTAAATTCTACTTTTATAGTTAGTCTTTTTTTTAGTTTTAAAACACCAAGAACTTAGTTTCGAATAAACACACATAAACAAACAAA |
| UAS349-FBP1h | TTAGTCAAAAAATTAGCCTTTTAATTCTGCTGTAACCCGTACATGCCCAAAATAGGGGGCGGGTTACACAGAATATATAACATCGTAGGTGTCTGGGTGAACAGTTTATTCCT**CCGGACGGATGGAATCGCCG**GAGCCCGCTTTTTA**CCGGACGGATGGAATCGCCG**AAGAATCCCAGCACCAAAATATTGTTTTCTTCACTAATAAGTATATAAAGACGGTAGGTATTGATTGTAATTCTGTAAATCTATTTCTTAAACTTCTTAAATTCTACTTTTATAGTTAGTCTTTTTTTTAGTTTTAAAACACCAAGAACTTAGTTTCGAATAAACACACATAAACAAACAAA |
| UAS349-Gis1 | TTAGTCAAAAAATTAGCCTTTTAATTCTGCTGTAACCCGTACATGCCCAAAATAGGGGGCGGGTTACACAGAATATATAACATCGTAGGTGTCTGGGTGAACAGTTTATTCCT**TAGGGAT**ACTAAATATAATGGAGCCCGCTTTTTAAGCT**TAGGGAT**AGAAAAAAAAAGAATCCCAGCACCAAAATATTGTTTTCTTCACTAATAAGTATATAAAGACGGTAGGTATTGATTGTAATTCTGTAAATCTATTTCTTAAACTTCTTAAATTCTACTTTTATAGTTAGTCTTTTTTTTAGTTTTAAAACACCAAGAACTTAGTTTCGAATAAACACACATAAACAAACAAA |
| UAS349-Gcr1D | TTAGTCAAAAAATTAGCCTTTTAATTCTGCTGTAACCCGTACATGCCCAAAATAGGGGGCGGGTTACACAGAATATATAACATCGTAGGTGTCTGGGTGAACAGTTTATTCCTTAAAGATACTAAATATAATGGAGCCCGCTTTTTAAGCTTAGAGATAGAAAAAAAAGAACACCCATACATCAAATATTGTTTTCTTCACTAATAAGTATATAAAGACGGTAGGTATTGATTGTAATTCTGTAAATCTATTTCTTAAACTTCTTAAATTCTACTTTTATAGTTAGTCTTTTTTTTAGTTTTAAAACACCAAGAACTTAGTTTCGAATAAACACACATAAACAAACAAA |
| TPI1pr | ATTTAAACTGTGAGGACCTTAATACATTCAGACACTTCTGCGGTATCACCCTACTTATTCCCTTCGAGATTATATCTAGGAACCCATCAGGTTGGTGGAAGATTACCCGTTCTAAGACTTTTCAGCTTCCTCTATTGATGTTACACCTGGACACCCCTTTTCTGGCATCCAGTTTTTAATCTTCAGTGGCATGTGAGATTCTCCGAAATTAATTAAAGCAATCACACAATTCTCTCGGATACCACCTCGGTTGAAACTGACAGGTGGTTTGTTACGCATGCTAATGCAAAGGAGCCTATATACCTTTGGCTCGGCTGCTGTAACAGGGAATATAAAGGGCAGCATAATTTAGGAGTTTAGTGAACTTGCAACATTTACTATTTTCCCTTCTTACGTAAATATTTTTCTTTTTAATTCTAAATCAATCTTTTTCAATTTTTTGTTTGTATTCTTTTCTTGCTTAAATCTATAACTACAAAAAACACATACATAAACTAAAA |
| TPI1-FBP1h | ATTTAAACTGTGAGGACCTTAATACATTCAGACACTTCTGCGGTATCACCCTACTTATTCCCTTCGAGATTATATCTAGGAACCCATCAGGTTGGTGGAAGATTACCCGTTCTAAGA**CCGGACGGATGGAATCGCCG**ATGTTACACCTGGACACCCCTTTTCTG**CCGGACGGATGGAATCGCCG**AGTGGCATGTGAGATTCTCCGAAATTAATTAAAGCAATCACACAATTCTCTCGGATACCACCTCGGTTGAAACTGACAGGTGGTTTGTTACGCATGCTAATGCAAAGGAGCCTATATACCTTTGGCTCGGCTGCTGTAACAGGGAATATAAAGGGCAGCATAATTTAGGAGTTTAGTGAACTTGCAACATTTACTATTTTCCCTTCTTACGTAAATATTTTTCTTTTTAATTCTAAATCAATCTTTTTCAATTTTTTGTTTGTATTCTTTTCTTGCTTAAATCTATAACTACAAAAAACACATACATAAACTAAAA |
| TPI1-Gcr1D | ATTTAAACTGTGAGGACCTTAATACATTCAGACACTTCTGCGGTATCACCCTACTTATTCCCTTCGAGATTATATCTAGGAACCCATCAGGTTGGTGGAAGATTACCCGTTCTAAGACTTTTTACAGAAGTCTATTGATGTTACACCTGGACACCCCTTTTCTACAGAAGAGTTTTTAATCTTCAGTGGCATGTGAGATTCTCCGAAATTAATTAAAGCAATCACACAATTCTCTCGGATACCACCTCGGTTGAAACTGACAGGTGGTTTGTTACGCATGCTAATGCAAAGGAGCCTATATACCTTTGGCTCGGCTGCTGTAACAGGGAATATAAAGGGCAGCATAATTTAGGAGTTTAGTGAACTTGCAACATTTACTATTTTCCCTTCTTACGTAAATATTTTTCTTTTTAATTCTAAATCAATCTTTTTCAATTTTTTGTTTGTATTCTTTTCTTGCTTAAATCTATAACTACAAAAAACACATACATAAACTAAAA |
| TEF1pr | CACACACCATAGCTTCAAAATGTTTCTACTCCTTTTTTACTCTTCCAGATTTTCTCGGACTCCGCGCATCGCCGTACCACTTCAAAACACCCAAGCACAGCATACTAAATTTCCCCTCTTTCTTCCTCTAGGGTGTCGTTAATTACCCGTACTAAAGGTTTGGAAAAGAAAAAAGAGACCGCCTCGTTTCTTTTTCTTCGTCGAAAAAGGCAATAAAAATTTTTATCACGTTTCTTTTTCTTGAAAATTTTTTTTTTTGATTTTTTTCTCTTTCGATGACCTCCCATTGATATTTAAGTTAATAAACGGTCTTCAATTTCTCAAGTTTCAGTTTCATTTTTCTTGTTCTATTACAACTTTTTTTACTTCTTGCTCATTAGAAAGAAAGCATAGCAATCTAATCTAAGTTTTAATTACAAA |
| PGK1pr | AGAAGTACCTTCAAAGAATGGGGTCTTATCTTGTTTTGCAAGTACCACTGAGCAGGATAATAATAGAAATGATAATATACTATAGTAGAGATAACGTCGATGACTTCCCATACTGTAATTGCTTTTAGTTGTGTATTTTTAGTGTGCAAGTTTCTGTAAATCGATTAATTTTTTTTTCTTTCCTCTTTTTATTAACCTTAATTTTTATTTTAGATTCCTGACTTCAACTCAAGACGCACAGATATTATAACATCTGCATAATAGGCATTTGCAAGAATTACTCGTGAGTAAGGAAAGAGTGAGGAACTATCGCATACCTGCATTTAAAGATGCCGATTTGGGCGCGAATCCTTTATTTTGGCTTCACCCTCATACTATTATCAGGGCCAGAAAAAGGAAGTGTTTCCCTCCTTCTTGAATTGATGTTACCCTCATAAAGCACGTGGCCTCTTATCGAGAAAGAAATTACCGTCGCTCGTGATTTGTTTGCAAAAAGAACAAAACTGAAAAAACCCAGACACGCTCGACTTCCTGTCTTCCTATTGATTGCAGCTTCCAATTTCGTCACACAACAAGGTCCTAGCGACGGCTCACAGGTTTTGTAACAAGCAATCGAAGGTTCTGGAATGGCGGGAAAGGGTTTAGTACCACATGCTATGATGCCCACTGTGATCTCCAGAGCAAAGTTCGTTCGATCGTACTGTTACTCTCTCTCTTTCAAACAGAATTGTCCGAATCGTGTGACAACAACAGCCTGTTCTCACACACTCTTTTCTTCTAACCAAGGGGGTGGTTTAGTTTAGTAGAACCTCGTGAAACTTACATTTACATATATATAAACTTGCATAAATTGGTCAATGCAAGAAATACATATTTGGTCTTTTCTAATTCGTAGTTTTTCAAGTTCTTAGATGCTTTCTTTTTCTCTTTTTTACAGATCATCAAGGAAGTAATTATCTACTTTTTACAACAAATATAAAACA |
| ADH2pr | TCTCTCCGGTTACAGCCTGTGTAACTGATTAATCCTGCCTTTCTAATCACCATTCTAATGTTTTAATTAAGGGATTTTGTCTTCATTAACGGCTTTCGCTCATAAAAATGTTATGACGTTTTGCCCGCAGGCGGGAAACCATCCACTTCACGAGACTGATCTCCTCTGCCGGAACACCGGGCATCTCCAACTTATAAGTTGGAGAAATAAGAGAATTTCAGATTGAGAGAATGAAAAAAAAAAAAAAAAAAAAGGCAGAGGAGAGCATAGAAATGGGGTTCACTTTTTGGTAAAGCTATAGCATGCCTATCACATATAAATAGAGTGCCAGTAGCGACTTTTTTCACACTCGAAATACTCTTACTACTGCTCTCTTGTTGTTTTTATCACTTCTTGTTTCTTCTTGGTAAATAGAATATCAAGCTACAAAAAGCATACAATCAACTATCAACTATTAACTATATCGTAATACACA |
